# Proliferative Exhausted CD8 T Cells Exacerbate Long-lasting Anti-tumor Effects in Human Papillomavirus Positive Head and Neck Squamous Cell Carcinoma

**DOI:** 10.1101/2022.10.27.513993

**Authors:** Danni Cheng, Ke Qiu, Yufang Rao, Minzi Mao, Li Li, Yan Wang, Yao Song, Junren Chen, Xiaowei Yi, Xiuli Shao, Shao Hui Huang, Yi Zhang, Xuemei Chen, Sisi Wu, Shuaishuai Yu, Jun Liu, Haiyang Wang, Xingchen Peng, Daibo Li, Lin Yang, Li Chen, Zhiye Ying, Yongbo Zheng, Meijun Zheng, Binwu Ying, Xiaoxi Zeng, Wei Zhang, Wei Xu, Geoffrey Liu, Fei Chen, Haopeng Yu, Yu Zhao, Jianjun Ren

**Author notes:** These authors contributed equally to this work (Danni Cheng, Ke Qiu, Yufang Rao, Minzi Mao). Senior author (Geoffrey Liu). **Corresponding authors:** Jianjun Ren, Department of Oto-Rhino-Laryngology and West China Biomedical Big Data Center, West China Hospital, Sichuan University, Chengdu, China, Yu Zhao, Department of Oto-Rhino-Laryngology and West China Biomedical Big Data Center, West China Hospital, Sichuan University, Chengdu, China, Haopeng Yu, West China Biomedical Big Data Center, West China Hospital, Sichuan University, Chengdu, China, Fei Chen, Department of Oto-Rhino-Laryngology, West China Hospital, Sichuan University, Chengdu, China. **Funding** National Natural Youth Science Foundation of China grant 82002868 (RJJ) National Natural Youth Science Foundation of China grant 32100927 (YHP) China Postdoctoral Science Foundation grant 2020M673250 (RJJ) The Science and Technology Department of Sichuan Province grant 2022YFS0066 (ZYB) The Science and Technology Department of Sichuan Province grant 2020YFS0111 (RJJ) The Science and Technology Department of Sichuan Province grant 2021YFS0158 (YHP) West China Hospital, Sichuan University grant ZYJC21027 (ZY) West China Hospital, Sichuan University grant 2019HXBH079 (RJJ) Sichuan University grant 2020SCU12049 (RJJ) The Health Department of Sichuan Province grant 20PJ030 (RJJ).

## Abstract

The survival prognosis of human papillomavirus (HPV)-positive and HPV-negative head and neck squamous cell carcinoma (HNSCC) is largely different, and little is known about the anti-tumor mechanism of tumor-infiltrated CD8^+^ exhausted T cells (Tex) in HNSCC. We performed cell level multi-omics sequencing on human HNSCC samples to decipher the multi-dimensional characteristics of Tex cells. A proliferative exhausted CD8^+^ T cell cluster (P-Tex) which was beneficial to survival outcomes of patients with HPV-positive HNSCC was identified. Interestingly, P-Tex cells expressed CDK4 genes as high as cancer cells, which could be simultaneously inhibited by CDK4 inhibitors and might be a potential reason for the ineffectiveness of CDK4 inhibitors in treating HPV-positive HNSCC. P-Tex cells could aggregate in the antigen presenting cell niches and activate certain signaling pathways. Together, our findings suggest a promising role for P-Tex cells in the prognosis of patients with HPV-positive HNSCC by providing modest but persistent anti-tumor effects.

**Graphic abstract:** 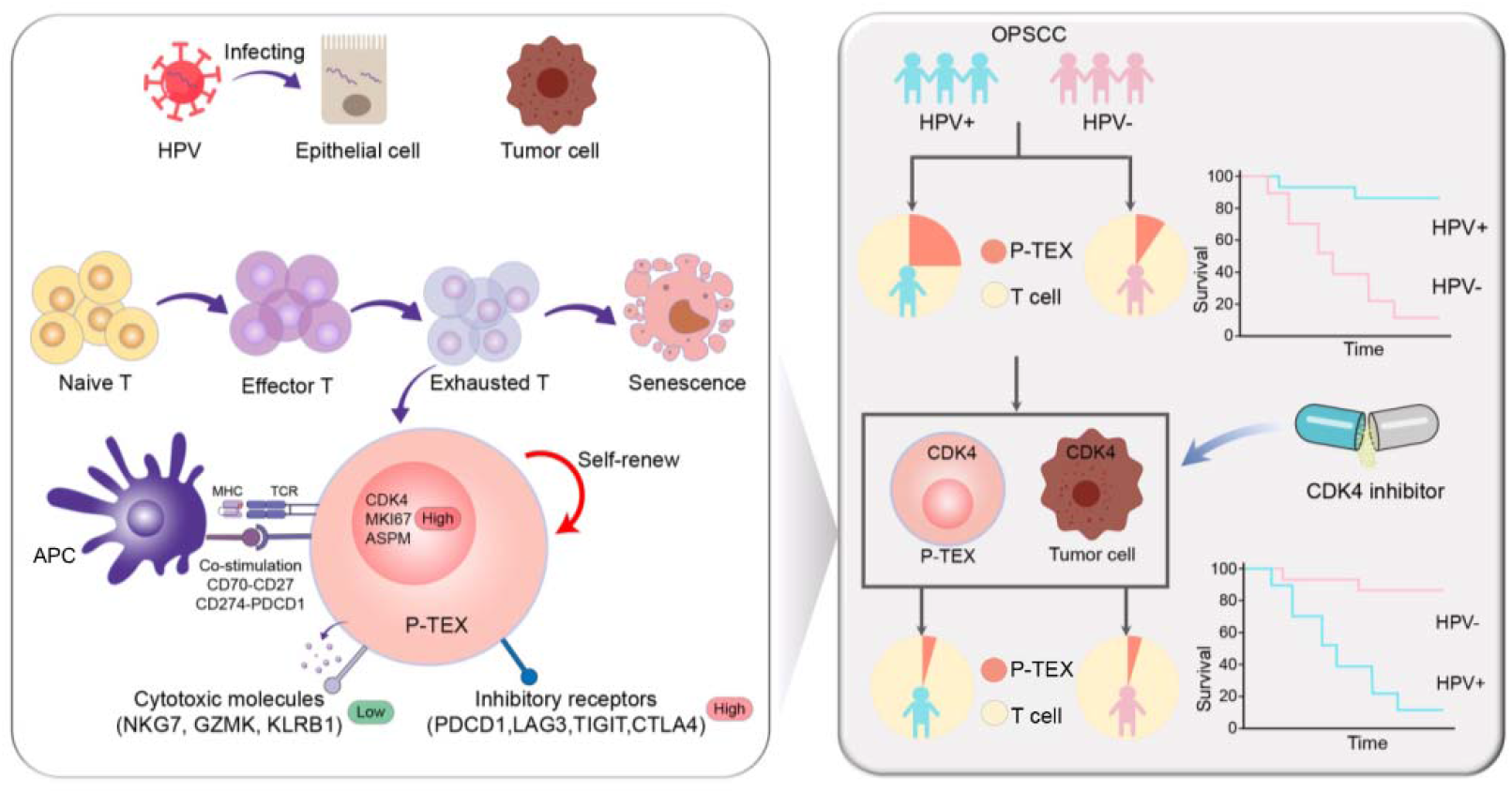

## Introduction

The incidence of head and neck squamous cell carcinoma (HNSCC) has continued to rise and an annual increase of 1.08 million new cases is estimated by 2030, which is greatly attributed to the increasing rates of human papillomavirus (HPV) infection(1–3). HPV-positive HNSCC and HPV-negative HNSCC displayed markedly different characteristics from pathogenesis to treatment outcomes and are visualized as two distinct clinical entities(4,5).

T cell exhaustion within the tumor microenvironment (TME) is a newly discovered phenomenon resulting from persistent antigen stimulation from chronic virus infection or tumors, in which tumor-infiltrated CD8^+^ T cells experience gradual alterations in their functional capacity while highly expressing multiple inhibitory receptors, including PD1, TIM3, LAG3, CTLA4 and TIGIT(6,7). Recent studies have revealed that heterogeneity is a hallmark of T cell exhaustion, and several distinct subsets of exhausted CD8^+^ T cells (Tex) have been identified, each with unique gene signatures, functional characteristics and epigenetic modifications(8,9). Meanwhile, specific transitions among those subsets have also been illustrated under certain circumstances and might be associated with retained effector function and enforced tumor control(10,11). However, how T cell exhaustion affects the prognosis of HNSCC patients and how it differs in HPV-positive and HPV-negative HNSCC remain to be further clarified.

Recently, there has been increasing evidences showing that HPV-positive HNSCC displays a T cell-inflamed phenotype distinct from its HPV-negative counterparts, indicating that HPV infection is associated with increased T cell infiltration and effector cell activation(12,13). Meanwhile, HPV infection has also been proven to be associated with T cell exhaustion, in which HPV-positive HNSCC expressed higher levels of multiple T-cell exhaustion markers such as PD1, TIM3, LAG3 and TIGIT compared to HPV-negative HNSCC, suggesting of stronger antigen-specific T cell immunity in HPV-positive HNSCC(12,14). However, more sophisticated T cell landscapes related to the HPV status of HNSCC remain to be further clarified.

Cyclin-dependent kinase 4 (CDK4) inhibitors are introduced as novel drugs by targeting and disrupting the CDK4-related cell cycle progression of cancer cells in recent years(15,16). However, promising treatment outcomes of CDK4 inhibitors were only observed in HPV-negative HNSCC rather than HPV-positive HNSCC(17–19) due to the mutation differences of cell cycle related genes in cancer cells, and less attention has been paid to the effects of CDK4 inhibitors on infiltrating T cells.

To address these questions, we applied cell-level multi-omics sequencing techniques to decipher the multi-dimensional characterization of tumor infiltrating T cells and its association with overall survival (OS) in human HNSCC with different HPV status.

## Results

### P-Tex cells were identified in the transcriptomic landscape of T cells in HNSCC patients

To decipher the multi-dimensional characterization of tumor infiltrating T cells in human HNSCC, we integrated multi-omics sequencing based on 5’ droplet-based single-cell RNA (scRNA-seq), single-cell TCR sequencing (scTCR-seq) and spatial transcriptomics in HNSCC samples (mainly oropharyngeal cell carcinoma, OPSCC, the most representative type of HPV-related HNSCC), and further verified in vitro (n=24, Table S1 and Figure. 1a). A total of 49,813 CD3^+^ T cells in 14 paired HNSCC tumor and normal adjacent samples were obtained after quality control. And 11 T cell clusters with distinct gene signatures were defined (Fig. 1b, Table S2), including four CD8^+^ T cell clusters (C1, C4, C7 and C9), five CD4^+^ T cell clusters (C2: naïve CD4^+^ T cells, C3: regulatory T cells [Treg], C5: T helper cells, C8: CD4-Tex, and C10: T follicular helper cells [Tfh]), one γδ T cell cluster (C6) and one double negative T cell cluster (C0: DN, CD4^-^CD8^-^), with marker genes(20) shown in Table S3.

**Figure 1.**
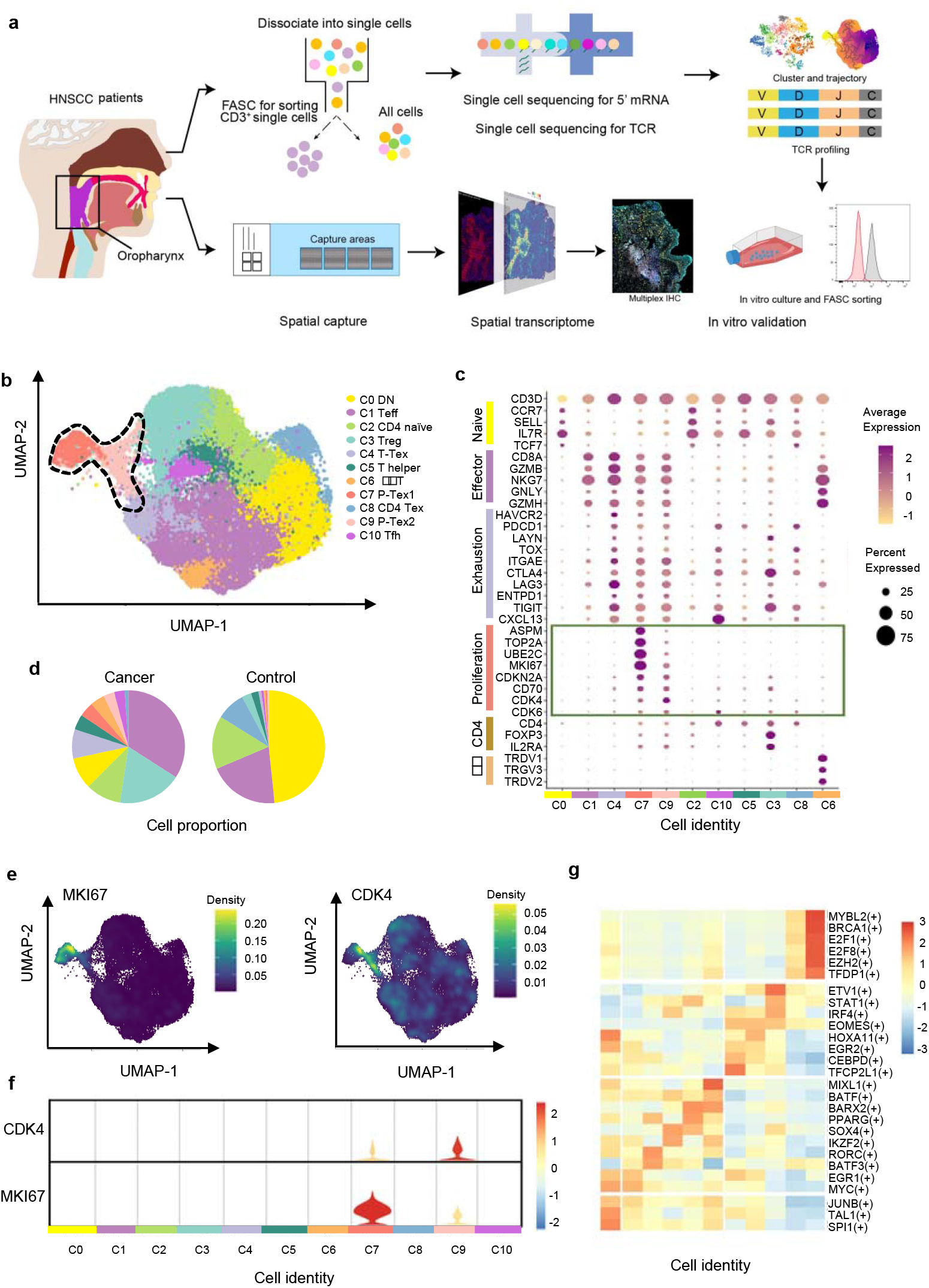
The P-Tex cell clusters identified in the T cell landscapes of HNSCC patients by scRNA-seq. **a,** The flow chart of this study. **b,** UMAP plot of all single T cells from 14 samples via 10× Genomics. Eleven T cell clusters with different functions are identified. **c,** Dotplot of selected T cell function-associated genes across different T cell clusters, showing both gene expression level (the color gradient) and the percentage of cells (the size of circle) in a given cluster. **d,** Pie charts of cell-type fractions identified in cancer and normal adjacent tissues, colored by cell types. **e,** The kernel density estimation plot showing the distribution of MKI67 and CDK4 genes of T cells. **f,** The gene expression levels of CDK4 and MKI67 shown by violin plots. **g,** Heatmap of the transcriptional regulators of top expressed genes in each T cell clusters.

We specifically characterized marker genes of CD8^+^ T cell into several panels based on their canonical biological function (proliferation, exhaustion and cytotoxicity, shown in Fig. 1c). Among the four CD8^+^ T cell clusters, C1 was characterized by expressing multiple effector genes, including *NKG7, GZMH, IFNG* and *KLRG1*, with a high cytotoxicity score but a low exhaustion score, thus representing effective CD8^+^ T cell (Teff) (21). C4 showed high expression of checkpoint marker genes, including *PDCD1, HAVCR2, LAG3*, and *TNFRSF9*, with both high cytotoxicity and exhaustion scores, which was consistent with the cell identity of terminally differentiated exhausted CD8^+^ T cells (T-Tex)(6). Notably, in addition to high expression of the above-mentioned checkpoint marker genes, C7 and C9 also displayed high expression levels of cell cycle-related genes, including *CCNA2, UBE2C* and *CDK4*, as well as stem-like genes *MKI67* (marker gene of proliferation) and *ASPM* (involved in regulation of the mitotic spindle and coordination of mitotic processes), with high cytotoxicity, exhaustion and proliferation genes, featuring a gene expression profile reminiscent of a previously reported Tex subset with high proliferative capacity(22), which we defined as P-Tex in the present study.

Meanwhile, T cells appeared to exhibit distinct tissue distributions, with higher proportions of Treg, Teff, T-Tex and P-Tex cells being observed in tumor tissues while more DN and CD4-Tex cells being observed in adjacent normal tissues (Fig. 1d, Supplementary Fig. 1a). To evaluate individual heterogeneity, we further clustered the cells of each HNSCC sample and confirmed the existence of all cell clusters across all samples (Supplementary Fig. 1b-c).

Notably, the P-Tex cluster (~2819 cells) could be further partitioned into two sub-clusters, which we annotated as P-Tex1 and P-Tex2. More specifically, *CDK4* (a canonical cell cycle-related marker gene) was mainly expressed in P-Tex2, whereas *MKI67* (the most canonical marker gene for proliferation) was mainly expressed in P-Tex1 (Fig. 1e-f), indicating that these two cell clusters might fulfill their proliferative capacity through different mechanisms. Moreover, to further investigate the potential upstream regulatory mechanisms in shaping the molecular characteristics of each unique T cell cluster, we analyzed the transcription factor networks that driving the expression of the top expression genes in each T cell cluster (shown in Fig. 1g and Table S4). Specifically, MYBL2, BRCA1, E2F1, E2F8, EZH2 and TFDP1 were the identified upstream regulatory transcription factors that predominantly drove the expression of proliferation-related genes of P-Tex1 and P-Tex2.

Taken together, our clustering strategy generated 11 distinct T cell clusters in HNSCC, among which P-Tex cells with both exhausted and proliferative phenotypes were identified.

**Extended Figure 1.**
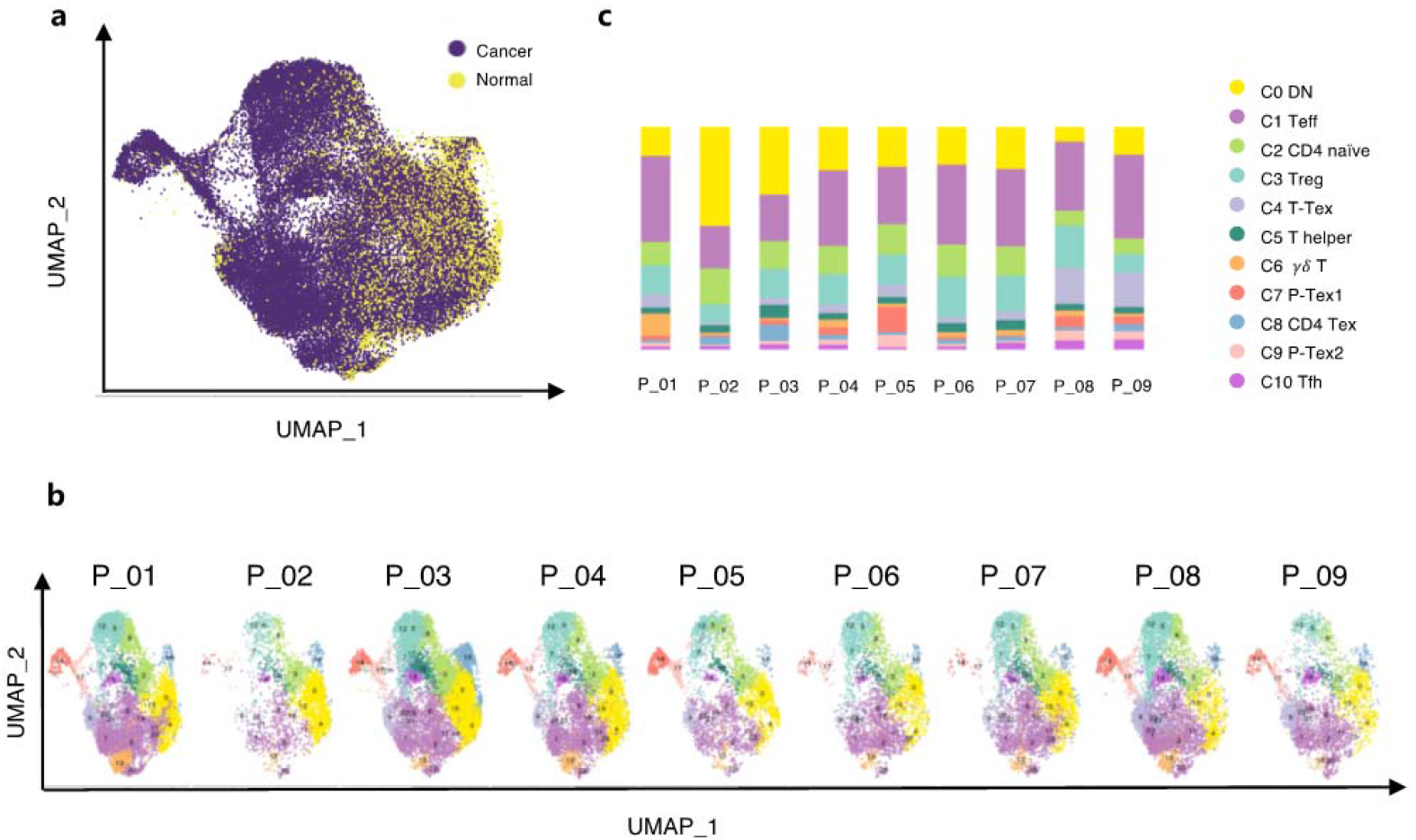
The extended summary of functional properties of T cell clusters in Figure 1. **a,** The distribution of T cells in cancer and adjacent normal tissues in UMAP plots, b, The distribution of T cells in each patient in UMAP plots, **c,** The distribution of cell proportions in each sample.

### Functional characteristics of P-Texs

To further investigate the functional characteristics of P-Tex cells, we characterized the function of marker genes by comparing with Teff and T-Tex clusters. Specifically, Gene Ontology (GO) enrichment analysis showed that T cell activation, lymphocyte differentiation and viral gene expression were enriched in all three Tex cell clusters, whereas the regulation of the cell cycle, apoptosis and certain immune responses were enriched in P-Texs, showing divergent functional specialization (Fig. 2a). Meanwhile, The activation states of CD8 T cells were compared by evaluating paired sc-RNA Seq and sc-TCR seq data, which reflects the magnitude of TCR signaling driving the differentiation of activated T cells into specific T cell subtypes(23). Our results showed that the T-Tex cluster was the most activated, followed by the two P-Tex clusters (Fig. 2b left). In addition, CD8 T cells in tumor tissues were more activated than those in adjacent normal tissues (Fig. 2b, right top). And no significant difference in T cell activation states was observed between HPV-positive and HPV-negative samples (Fig. 2b right bottom).

**Figure 2.**
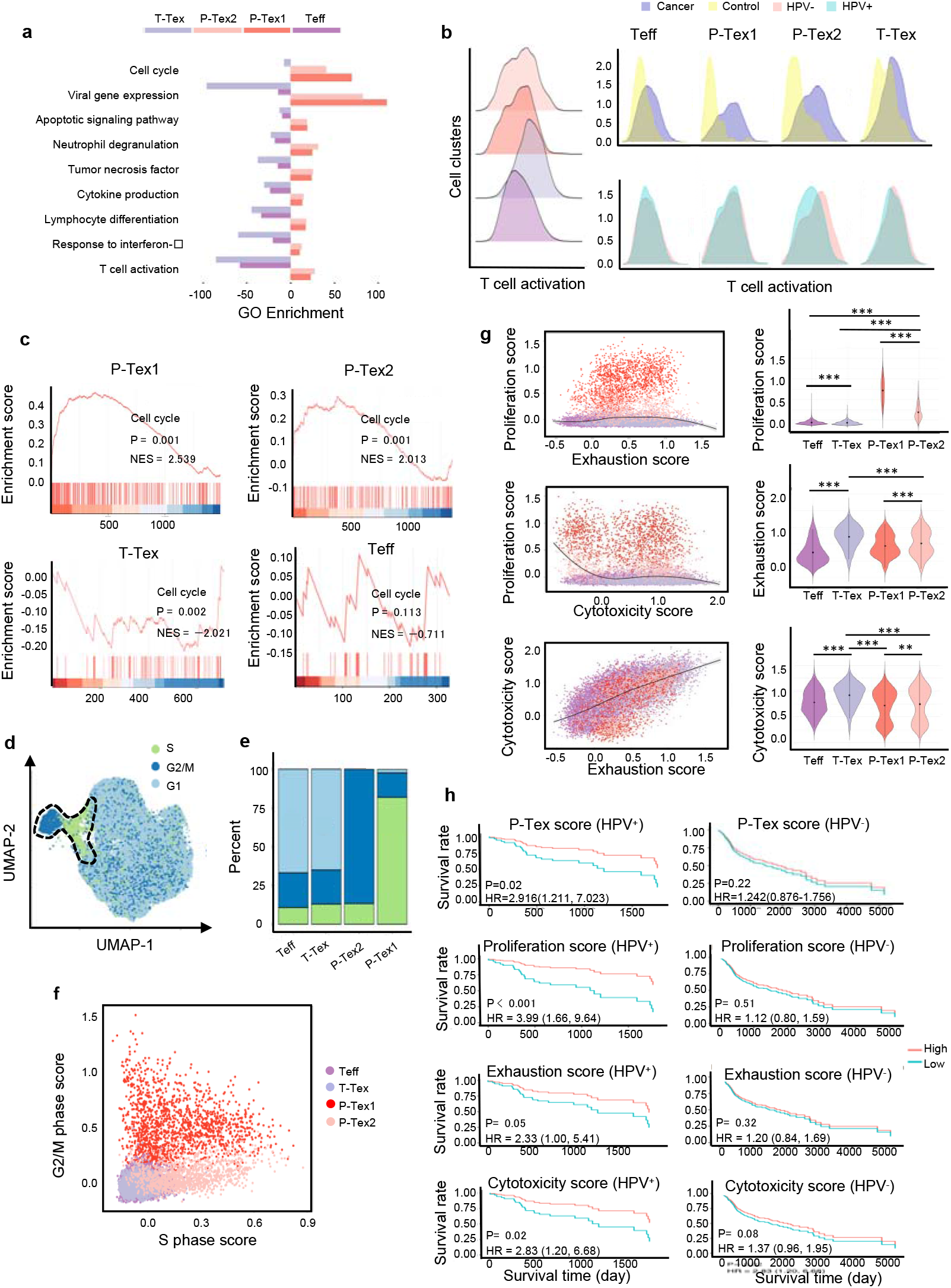
The comparison of functional characteristics between P-Texs and other CD8 T cell clusters. **a**, Gene ontology (GO) analysis of differentially expressed genes in each CD8 T cell clusters, colored by cell types. **b**, Histogram of activation states of all CD8 T cells (left) and CD8 T cells separated by tissue types (right top) or HPV status (right bottom) using paired single-Cell TCR Seq and RNA-Seq data. **c**, The GSEA diagrams show the enrichment profiles of cell cycle pathway in each CD8 T cell clusters. **d-f**, The distribution and scores of cell cycle phases of each CD8 T cells. **g**, The proliferation, exhaustion and cytotoxic scores of each CD8 T cell clusters. Proliferation score: averaged expression of MKI67 related genes; exhaustion score: averaged expression of PDCD1 related genes; cytotoxic score: averaged expression of GZMB related genes. **h**, The Kaplan–Meier curves show the overall survival rate of HPV^+^/HPV^-^ HNSCC patients with different proliferation, exhaustion and cytotoxic scores in TCGA cohort, adjusted for age and gender. ***: p<0.001, **: p<0.01, *: p<0.05.

Additionally, to further confirm that P-Texs displayed high cell cycle-related function, we performed Gene Set Enrichment Analysis (GSEA) using the gene set that represents the cell cycle pathway, and the results showed that two P-Tex clusters were more enriched in the cell cycle signal pathway than the T-Tex and Teff clusters (p<0.001, Fig. 2c). To further investigate the cell cycle phase of each T cell cluster, we calculated cell cycle scores and visualized them on UMAP plots (Fig 2d, Table S5). P-Tex1 was mainly in G2/M phase, indicating cells were under a proliferative burst, which was consistent with the high expression of proliferation marker gene *MKI67*(24), whereas P-Tex2 was mainly in S phase, an essential phase for DNA replication before undergoing mitosis, which was consistent with the high expression of *CDK4* (initiating the G1 to S phase transition) (Fig 2e-f)(24–26). Taken together, P-Tex1 and P-Tex2 might represent proliferation cells in two distinct cell cycle phases.

It is also noteworthy that HPV-positive HNSCC patients in TCGA (The Cancer Genome Atlas) cohort with higher P-Tex scores (5-year OS: 55.8% vs 22.6%, p=0.02), proliferation score (*MKI67*-related genes, 5-year OS: 49.1% vs 15.8%, p<0.001), exhaustion score (*PDCD1*-related genes, 5-year OS: 56.2% vs 23.1%, p=0.05), or cytotoxic score (*GZMB*-related genes, 5-year OS: 55.4% vs 21.9%, p=0.02) had better survival outcomes, whereas similar trends were not observed in HPV-negative HNSCC patients (Fig. 2g-h, Table S5) It is probably related to the difference in tumor microenvironment between HPV^+^ vs HPV^-^ HNSCC.

Taken together, the P-Tex cluster displayed high expression levels of proliferation- and cell cycle-related genes. More importantly, HPV^-^ positive HNSCC patients with higher P-Tex score, proliferation score, exhaustion score or cytotoxic score had better survival outcomes, while this trend was not observed in HPV-negative HNSCC patients.

### Paired scRNA-seq and TCR-seq Revealed the developmental trajectory of P-Tex

Given the clonal accumulation of CD8 T cells was a result of local T cell proliferation and activation in the tumor environment(27), we integrated paired scRNA-seq and TCR-seq data and performed pseudotime trajectory analysis to further quantitatively assess the activation states and to trace the lineage relationships of T cells(28). After quality control, we obtained TCRs with both alpha and beta chains for 33,897 T cells, including 20,607 unique TCRs, 2,798 double TCRs and 10,492 clonally expanded TCRs, with clonal sizes ranging from 2 to 162.

To further confirm whether the T cell clonality was associated with TME and HPV status, we systematically compared the clonality by cell clusters, tissue origin and HPV status, respectively. The CD8^+^ T cell clusters harbored more clonally expanded cells than CD4^+^ T cell clusters and DN cell clusters in general, among which Tex harbored the highest proportions of clonal cells, followed by the two P-Tex clusters, which were more abundant than the Teff cells (Fig. 3a-b, Supplementary Fig. 2a-b). Our results showed that hyperexpanded TCR clonotypes were more enriched in tumors than adjacent normal tissues (Fig. 3c-d, Supplementary Fig. 2c-d). However, the proportions of hyperexpanded TCR clonotypes of Teff, P-Tex1 and T-Tex showed no significant difference between HPV-positive samples and HPV-negative samples (Fig. 3e-f, Supplementary Fig. 2e-f). Correspondingly, a higher diversity of TCRs was observed in adjacent normal tissues and HPV-negative samples (Fig. 3g), indicating the absence of a strong antigen-specific immune response, which further confirmed the crucial roles of virus and tumor play in local T cell proliferation and activation(29,30).

**Figure 3.**
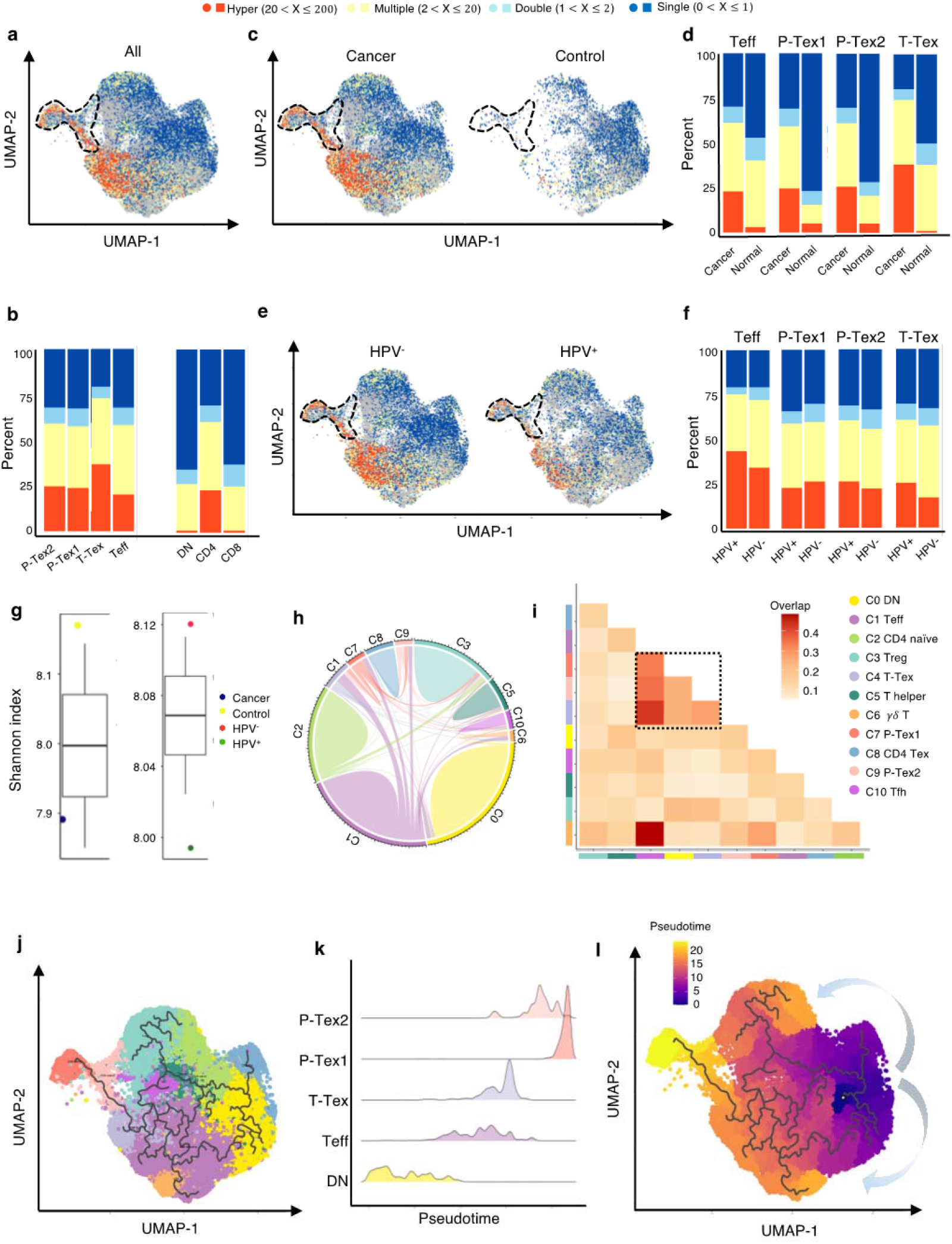
The developmental trajectory and lineage relationships among T cell clusters. **a-f,** Single-cell TCR protiling of HNSCC in all samples **(a-b),** cancer tissues vs. normal tissues **(c-d),** HPV^+^ *vs*. HPV^-^ **(e-f).** Bar plots show the fractions of each clonotype frequencies. The clonotype frequencies are defined as unique (π= 1), double (n=2), multiple clones (2<n≤20) and hyper clones (20<n≤200) according to the numbers of clonotypes. **g**. The TCR diversity of cancer tissues *vs*. normal tissues and HPV^+^ *vs*. HPV^-^ samples, calculated using Shannon metric, **h-i,** Cell state transition of T cell clusters inferred by shared TCRs. The chord Diagram (**h**) showing the fraction of shared clonotypes among each cell clusters. Lines connecting different clusters are based on the degree of TCR sharing, with the width of lines representing the number of shared TCRs. The cional overlap diagram (**i**) measures the clonal similarity among each cluster. Color gradient in the grid reters to the overlap coefficient. The higher the index score, the higher the clonal diversity, **j-l.** Potential developmental trajectory of T cells inferred by Monocle3 based on gene expressions.

We further examined the TCR clonotype occupation among each cluster and revealed that most of the shared TCRs were observed among the T-Tex, P-Tex and Teff clusters (Fig. 3h-i, Table S6-7). The Teff cluster had higher proportion of TCRs shared with the γδ T (overlap coefficient, oc=0.49), Tex (oc=0.43), P-Tex2 (oc=0.35) and P-Tex1 (oc=0.33) clusters, respectively, indicating they had common ancestry of origin. Besides, Supplementary Fig. 2g-j and Supplementary Fig. 3a-c shows the distribution of the top shared clonotypes across CD8^+^ T cell clusters, individuals and HPV status. There was almost no shared TCRs across individuals, indicating the highly heterogeneous characteristics of T cells among individuals.

To further investigate their lineage relationships, we performed pseudotime analysis for CD3^+^ T cells on the basis of transcriptional similarities (Fig. 3j-l, Supplementary Fig. 3d). The starting point of pseudotime was the DN cluster, with CD4^+^ T cell and CD8^+^ T cell clusters differentiating toward two different directions, suggesting of their distinct developmental paths. Notably, two P-tex clusters primarily aggregated at the end of the pseudotime backbone of CD8^+^ T cells, and presented to be a specific branch originating from T-Tex, demonstrating its specific activation state and characteristic, which was distinct from other T-Tex cells. Besides, P-Tex2 was located ahead of P-Tex1 on the pseudotime, which was consistent with the results in Fig. 2d-e, where P-Tex2 cells mainly entered the S phase of the cell cycle (early phase), while P-Tex1 mainly entered the G2/M phase (later phase).

Taken together, given that two P-Tex clusters were located at the end of the developmental trajectory of the Teff and T-Tex cells and that P-Tex clusters partially shared TCRs with the T-Tex cluster, we speculated that Teff cells transformed from an activated to exhausted state (T-Tex cells), and some of the T-Tex cells could further gradually transform into a unique P-Tex subpopulation with a highly specific proliferation state.

**Extended Figure 2.**
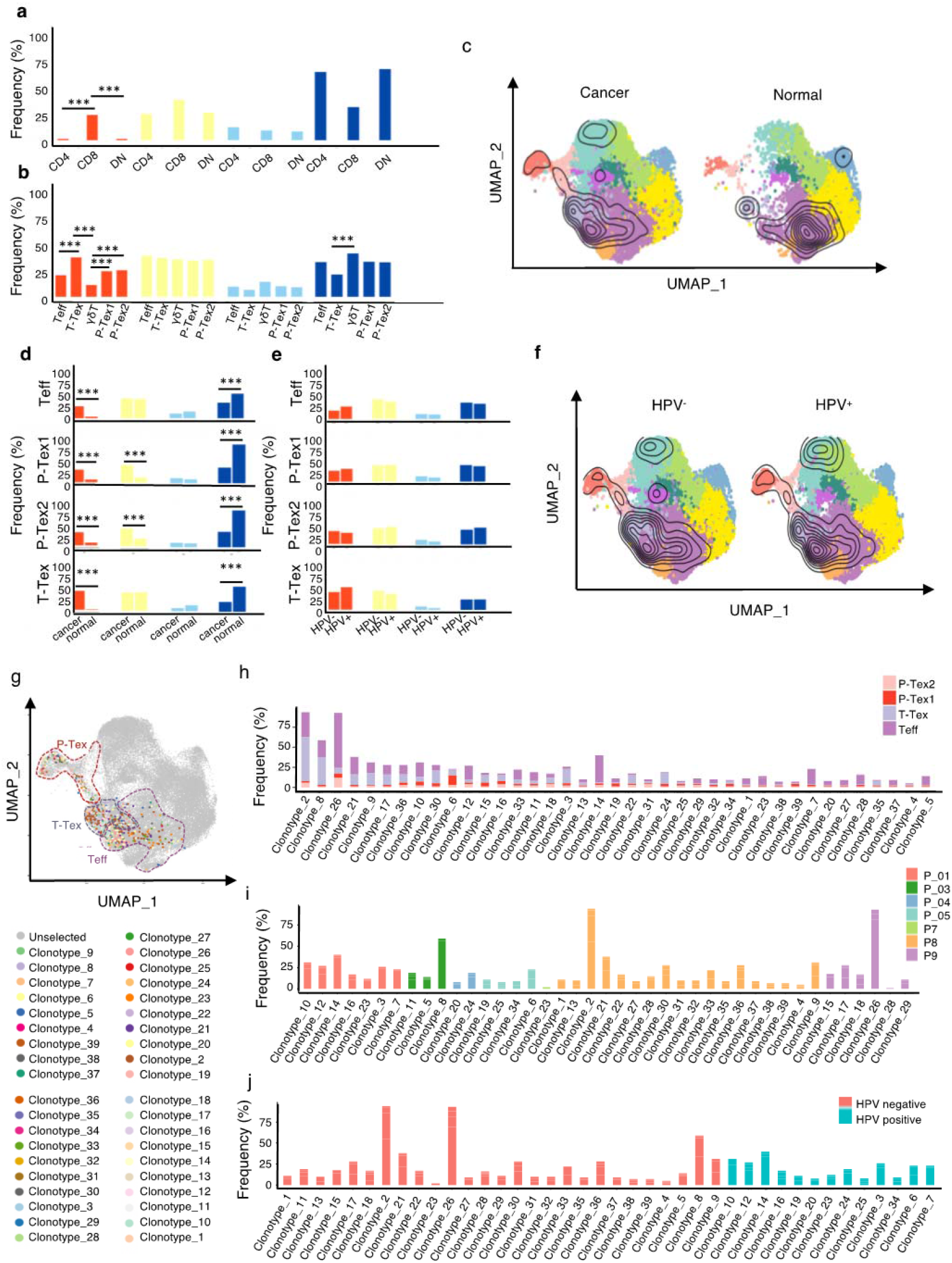
Extended summary of TCR properties of Figure 3. **a-b,** The supplementary comparison of the clonotype frequencies of clusters in each cluster **(a-b),** cancer tissues *vs*. normal tissues **(c-d),** HPV^+^ *vs*. HPV^-^ **(e-f).** The clonotypes are defined as unique (n =1), double (n=2), multiple (2<n≤20), and hyper clonal (20<n≤200) according to their clonotype numbers. The clonalOverlay diagrams show the clonal expansion in HPV^+^ *vs*. HPV^-^ and cancer tissues vs. normal tissues by overlaying the cells with specific clonal frequency onto the UMAP plots in Seurat **(c and f).** The density contours indicate the frequencies of TCR, with the number of clones ≥ 3 to be the cut-off value of the outermost circle layer, and the most central circle layer represents the area with the highest TCR expansion, **g-j**. The distribution of shared CD8+ clones on the UMAP plot. Colored dots were CD8+ cells of identical clonolypes. The colored circles highlight the cluster information of each cell, as defined in Figure 1a, with fractional Teff, T-Tex and P-Tex cells sharing the same TCRs shown in colors. Bar charts show the proportion of the each clonotype in each cluster (**h**), samples (**i**) and different **HPV** status (**j**), respectively.

### The self-renewal, proliferation capacity and cytotoxicity of P-Texs in vitro

To further verify the aforementioned function of P-Tex cells, we sorted P-Tex cells (CD3^+^CD8^+^UBE2C^+^PD1^+^) and T-Tex cells (CD3^+^CD8^+^UBE2C^-^KLRB1^+^PD1^+^) via flow cytometry to compare their functions in vitro (Fig. 4a-g). Unexpectedly, our flow cytometry results showed that the proliferation rate of P-Tex cells cultured with IL-2 for 15 days and 20 days in vitro was much slower than that of T-Tex cells (Fig. 4a), whereas the cell viability of P-Tex cells was much higher than that of T-Tex cells (Fig. 4b-c), indicating that instead of showing high proliferation capacity when stimulated in vitro, the P-Tex cells mainly maintained high self-renewal capacity.

**Figure 4.**
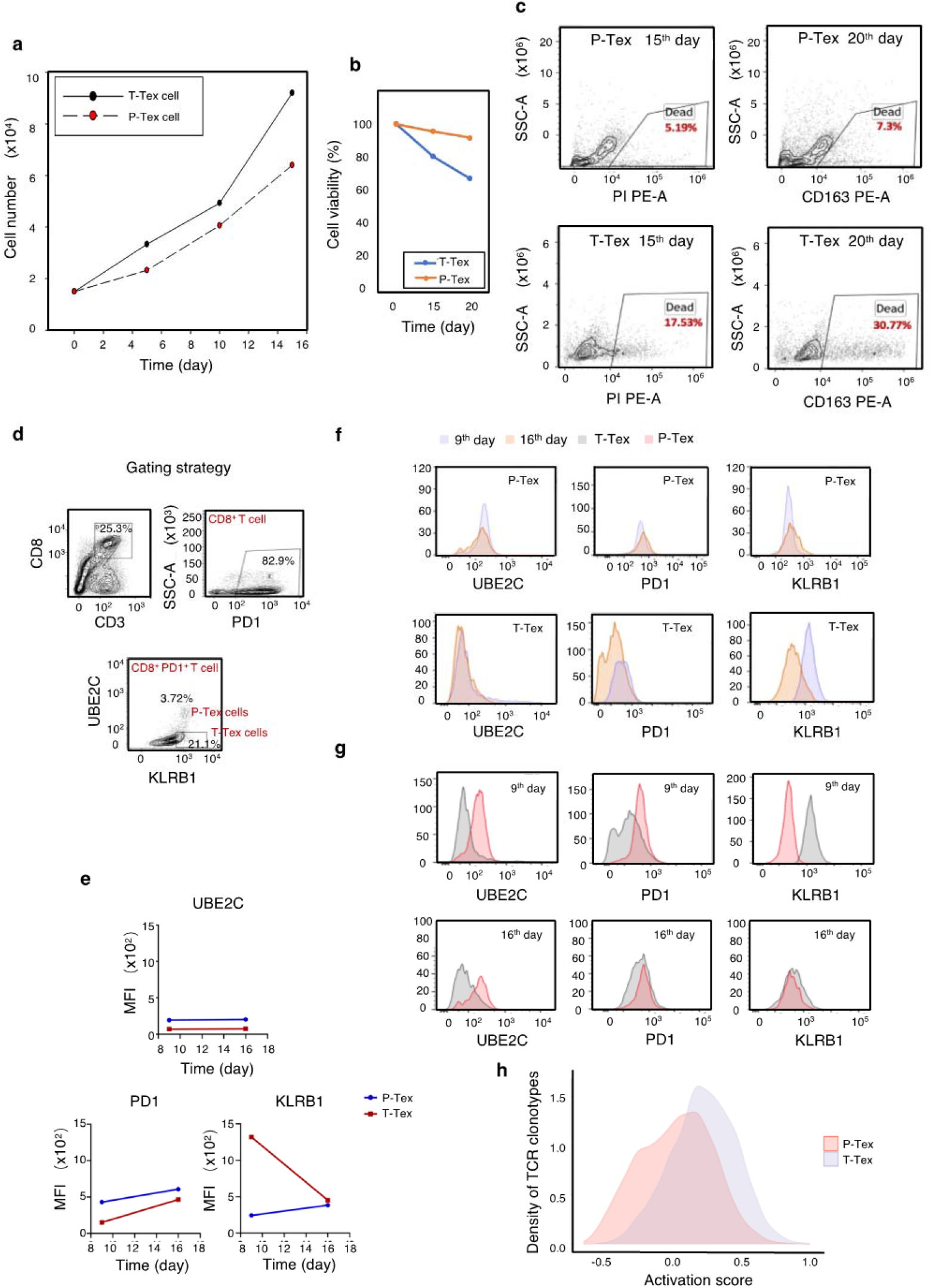
The self-renew and proliferation capacity of P-Texs in vitro. **a-c,** Comparing the proliferation (a) and self-renew (**b-c**) capacity of P-Tex and T-Tex cells cultured with IL-2 for 15 days and 20 days in vitro measured by CCK8 experiment, **d-g,** Representative flow cytometry assay of UBE2C, PD1, and KLRB1 of P-Tex cells after 9 and 15 days of stimulation with anti-CD3/CD28 microbeads in vitro, **d:** The gating strategies of PD1, KLRB1 and UBE2C. **e-g**: Mean Fluorescence Intensity (**e**) and cell count (**f-g**) of UBE2C, PD1, and KLRB1 in P-Tex and T-Tex cells at different days detected by flow cytometry, **h**, Histogram of activation states of P-Tex and Tex cells by paired Single-Cell TCR Sequencing and RNA-Seq data.

Moreover, P-Tex cells expressed higher levels of proliferation-related marker UBE2C, as measured by flow cytometry, and the variations of UBE2C in P-Tex and T-Tex cells between 9 days and 16 days were relatively stable (Fig. 4d-g). Besides, compared with T-Tex cells, P-Tex cells expressed higher exhaustion-related markers (PD1), and the expression of PD1 in P-Tex and T-Tex cells gradually increased from Day 9 to Day 16. Meanwhile, a larger proportion of T-Tex cells produced more cytotoxic-related markers (KLRB1) than P-Tex cells after stimulated with CD3/CD28 microbeads and IL-2 for 9 days and 16 days in vitro, whereas the expression of KLRB1 in T-Tex gradually decreased since Day 9, and the expression of KLRB1 in P-Tex was relatively stable. Meanwhile, the results of our in vitro experiments were consistent with the paired single-cell RNA-Seq and TCR-seq data showing that the activation states of T-Tex cells were higher than those of P-Tex cells (Fig. 4h).

Taken together, P-Tex cells represent a unique sub-cluster of the exhausted CD8 T cells, which maintain high self-renewal capacity in vitro and could provide modest but persistent anti-tumor effects.

### The effect of CDK4 inhibitor on P-Tex cells might be a reason for its ineffectiveness in HPV-positive HNSCC patients

To better understand the anti-tumor role of P-Tex within the TME, we additionally conducted 5’ droplet-based scRNA-seq profiles (10× Genomics) for primary tumors with paired adjacent normal tissues from two HNSCC patients. All biopsies were histologically examined by two independent pathologists. After quality control, a total of 13,515 cells from tumors (9,040 cells) and adjacent normal tissues (4,476 cells) were obtained. Given the fact that higher heterogeneity of cellular compositions exists in the TME than pure T cells, we recategorized all cells into 20 cell clusters according to previously reported markers (Fig. 5a, Supplementary Fig. 4a, Table S8). We consistently identified that P-Tex cells highly expressed proliferation- and cell cycle-related genes and functions as the cancer epithelial cluster (Fig. 5b, Supplementary Fig. 4b-d).

**Figure 5.**
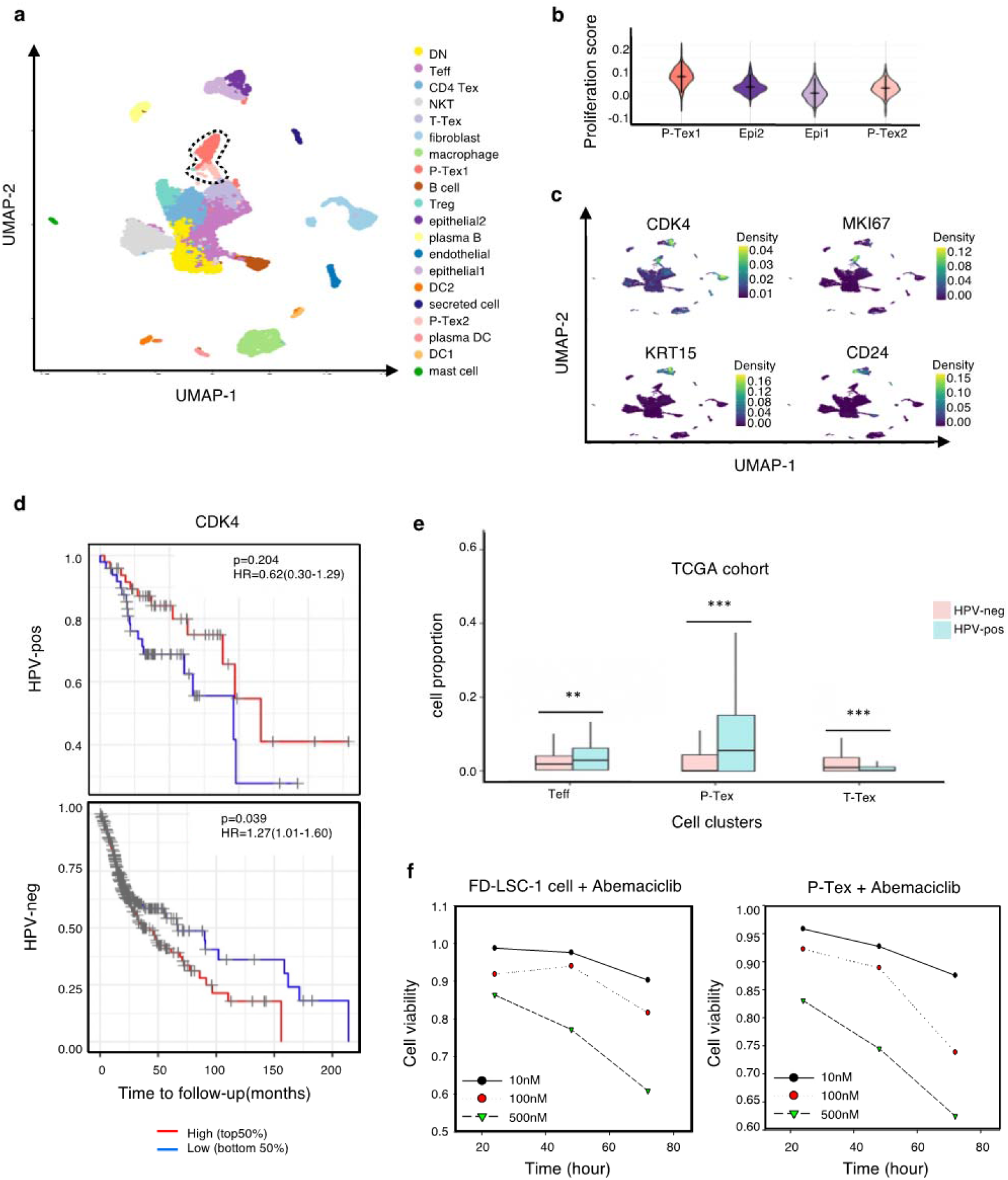
The expression of CDK4 gene in P-Tex2 cluster is associate with the treatment outcomes ot HPV^+^ HNSCC patients. **a,** Single-cell transcriptomic profiling of HNSCC TME. Twenty cell clusters are identified, colored by cell types, **b,** The proliferation status of P-Tex and epithelial cells in violin plot, **c**, The kernel density estimate distribution of proliferation markers (CDK4 and MKI67) and epithelial cancer cell markers (KRT15 and CD24) in UMAP plots, **d,** The overall survival rate of HPV^+^/HPV^-^ HNSCC patients in TCGA cohort related to the expression levels of CDK4 gene, adjusted for age and gender, **e,** The proportion of P-Texs, T-Tex and TEFF clusters in HPV^+^ and HPV^-^ samples in TCGA cohort by using the deconvolution algorithm. Marker genes that were used to define cell clusters in Figure 5a are deconvolved into the TCGA data to obtain the proportion of P-Texs, T-Tex and Teff clusters in the TCGA cohort, **f**, The cell viability of P-Tex and cancer epithelial cells assessed by CCK8 experiment after Abemaciclib treated in vitro.

P-Tex clusters were predominantly tumor-derived (Supplementary Fig. 5a-b) cells. We further investigated the expression and distribution of proliferation-related (*CDK4, MKI67*) and cancer-related epithelial (*KRT15, CD24*) cell marker genes among each cluster (Fig. 5c). Notably, *CDK4* was highly expressed in P-Tex2 cells, cancer epithelial cells and fibroblasts, while *MKI67* was highly expressed in P-Tex1 clusters. CDK4 is a well-known cancer treatment target, and CDK4 inhibitors (e.g., abemaciclib and palbociclib) have demonstrated cytostatic activity in HPV-negative HNSCC, whereas their effects on HPV-positive HNSCC are not obvious(17,18,31). And it was interesting that HPV-positive HNSCC patients with higher *CDK4* expression levels showed better survival than patients with lower *CDK4* expression, whereas HPV-negative HNSCC patients with higher *CDK4* expression levels showed worse prognosis (Fig. 5d). Besides, compared with HPV-negative patients, the proportion of P-Tex was higher in the TME of HPV-positive HNSCC patients (TCGA cohort, Fig. 5e, Supplementary Fig. 5c, Table S9). These findings raised the question of whether P-Tex cells that were beneficial to the prognosis of HPV-positive HNSCC patients would be simultaneously suppressed by CDK4 inhibitors.

To answer this question, we compared the cell viability of P-Tex cells and cancer epithelial cells by culturing with abemaciclib in vitro, respectively. As expected, abemaciclib inhibited the cell viability of both cancer cells (FD-LSC-1 cells) and P-Tex cells (Fig. 5f). Therefore, we speculated that the inhibition of CDK4 inhibitor on the cell viability of P-Tex cells (which were beneficial to the survival prognosis of HPV-positive HNSCC patients) might be a potential reason why CDK4 inhibitors were ineffective in treating HPV-positive HNSCC patients.

**Extended Figure 4.**
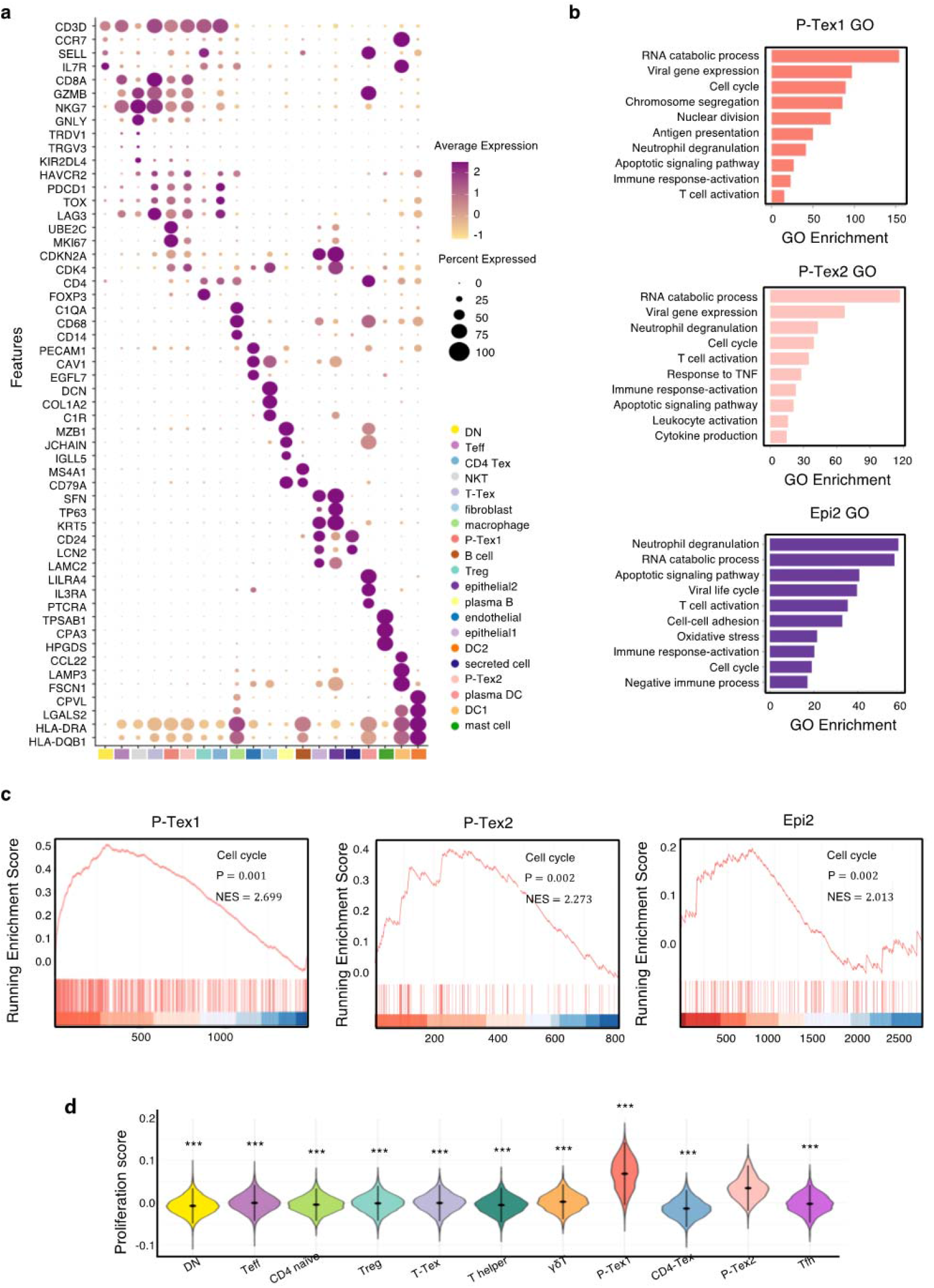
The extended summary of functional properties of cell clusters in Figure 5. **a**, Average expression of selected T cell function-associated genes across different ceil clusters, **b**, Gene ontology (GO) analysis of differentially expressed genes in two P-Tex and cancer epithelial cell clusters, **c,** The GSEA diagrams show the enrichment of cell cycle genes in P-Tex clusters and cancer epithelia cluster, **d,** The proliferation status of each T celi cluster.

#### The cell–cell interactions between T cells and APCs in the HNSCC TME

To determine the underlying mechanism by which P-Tex cells fulfill their proliferation-related and anti-tumor function, we systematically explored the crosstalk between T cells and other cells in the HNSCC tumor microenvironment (TME). The results showed that the interactions between P-Tex and Tex clusters and APCs (especially DCs) in the HNSCC TME were mainly enriched in T cell activation and proliferation signaling pathways, such as CD70-, CD80-, ICOS- and PD-L1-related signaling pathways (Fig. 6a-b, Supplementary Fig. 5d, Table S10). Given the fact that the colocalization of APCs and T cells is the precondition for fulfilling their function related to antigen presentation and T cell activation, we further conducted spatial transcriptome (ST) analysis for representative fresh HNSCC tumor (P_08) to verify their spatial distribution characteristics. We identified 17 spatial cluster areas, among which cluster 16 were defined as APC area (Fig. 6c). The P-Tex and Tex cells were characterized by the co-localization in the APC aggregation area with significantly higher P-Tex scores and Tex scores than other non-APC areas (P<0.001), and the correlations among the three scores in the APC area were higher than those in other non-APC areas. As expected, the activation score of T cells were higher in APC area (Fig. 6d-e, Supplementary Fig. 6, Table S11). Besides, the aforementioned ligand–receptor interactions of T cell activation and proliferation signaling pathways (CD70-CD27, CD80-ICOS, CD86-CTLA4, CD274-PDCD1) were also detected in the APC areas (Fig. 6f). We also observed enriched signaling pathways in APC areas involving the cell cycle, neutrophil activation and RNA splicing (Fig. 6g), supporting that these antigen-presenting cells play a role in modulating the immune response within the TME by promoting T-cell activation(32,33).

**Figure 6.**
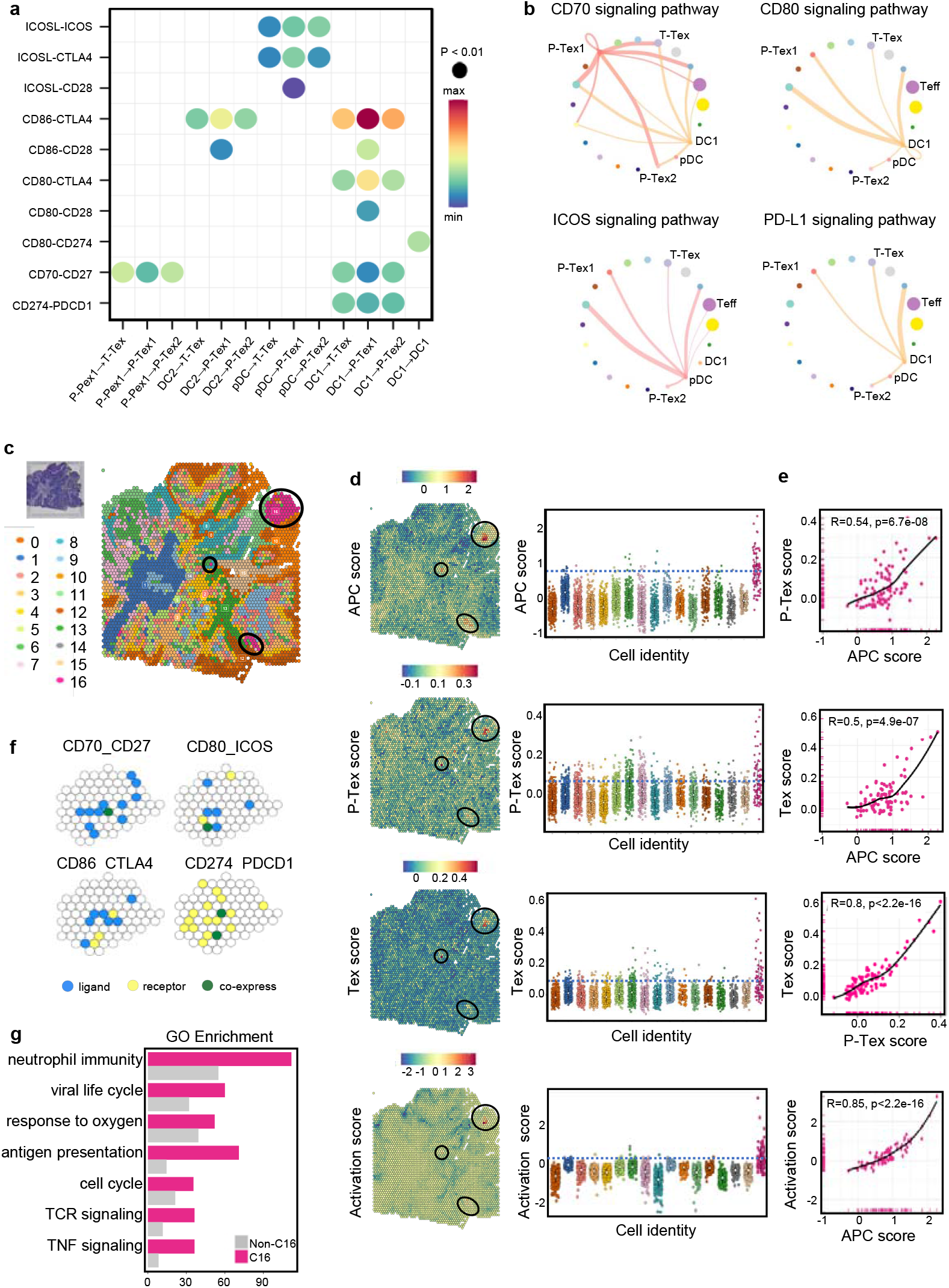
The cell–cell interactions between T cells and APC cells are enriched in the proliferation and cell activation pathways in HNSCC TME. **a**, The communication probabilities mediated by selected ligand–receptor pairs among different cell types. The color gradient shows the level of interaction. **b**, Network circle graphs visualize the inferred communication network of signaling pathways among different cell clusters derived by ligand–receptor interactions. The color of lines are consistent with the ligands. The width of lines are proportional to the interaction strength, and the circle sizes are proportional to the number of cells in each clusters. **c**, The spatial transcriptomic landscape of representative HNSCC samples. **d-e**, P-Tex and Tex features were co-expressed in APC area (cluster 16). The circles in SpatialDimPlot (**d**, left) represent APC, P-Tex and Tex scores enriched in the APC area (cluster 16). The Texs and P-Tex scores were higher in the APC aggregation area (**d**, right). The correlation of P-Tex, Tex, Activation scores and APC scores in the spatial transcriptome (**e**). **f**, Spatial feature plots of selected ligands–receptor interactions enriched in APC area. Spatial feature plots showing the expression pattern of single ligand genes (CD70, CD80, CD86, CD274, yellow spots), single receptor genes (CD27, ICOS, CTLA4, PDCD1, blue spots) and co-expression pattern (green spots) in APC area. **g**, GO analysis identified the enriched gene functions in APC area of spatial transcriptome.

**Extended Figure 5.**
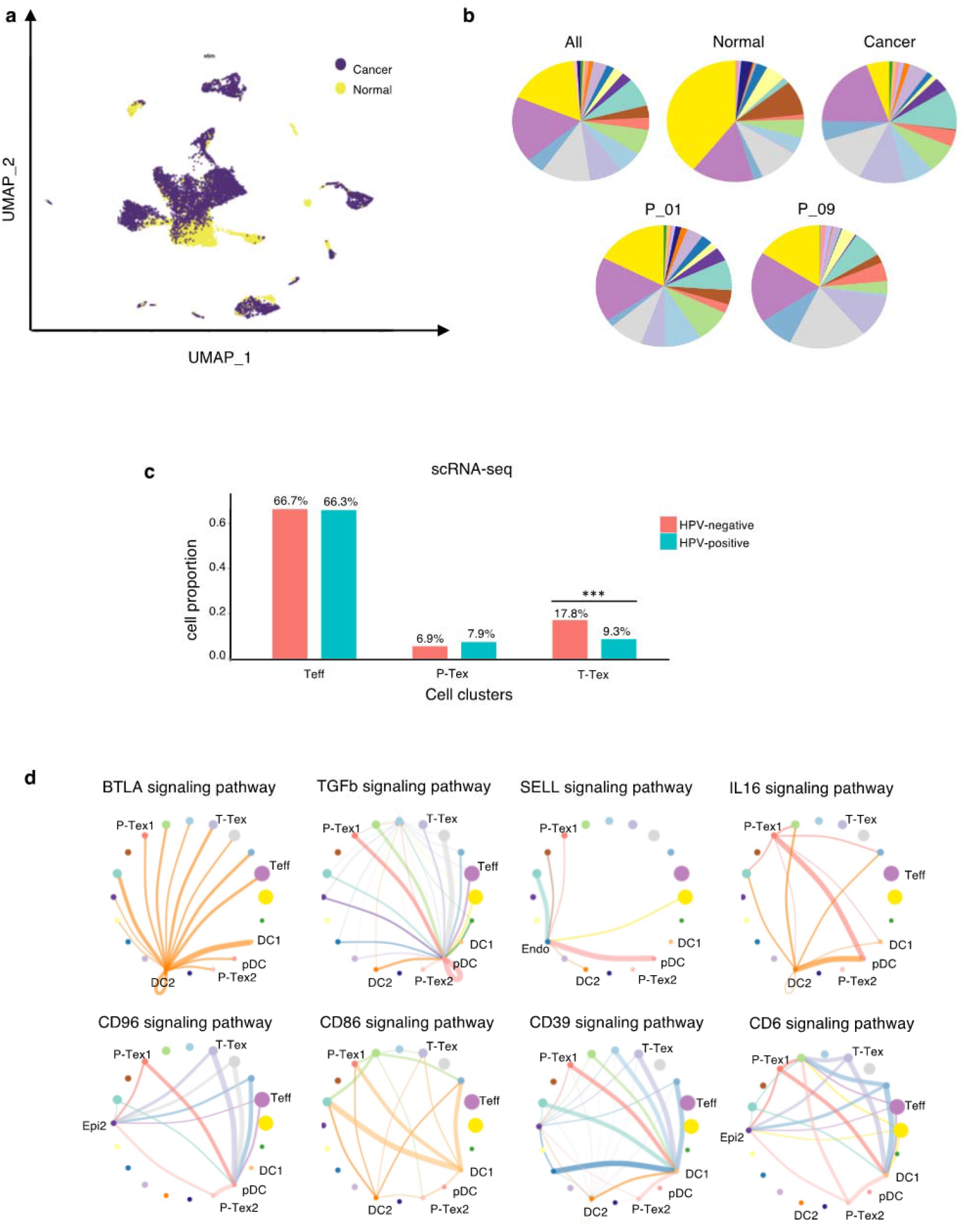
The extended summary of function properties of cell clusters in Figure 5. **a,** The distribution of cells in cancer and normal tissues in UMAP plots, **b,** The proportion of each cell cluster in all samples, cancer vs. normal tissue samples, and individual samples, colored by cell types, **c**, Comparing the proportion of P-Texs, T-Tex and Teff clusters in HPV+ and HPV− samples in single-cell sequencing data, **d**, The supplementary cell-cell interactions of HNSCC TME for Figure 6b.

**Extended Figure 6.**
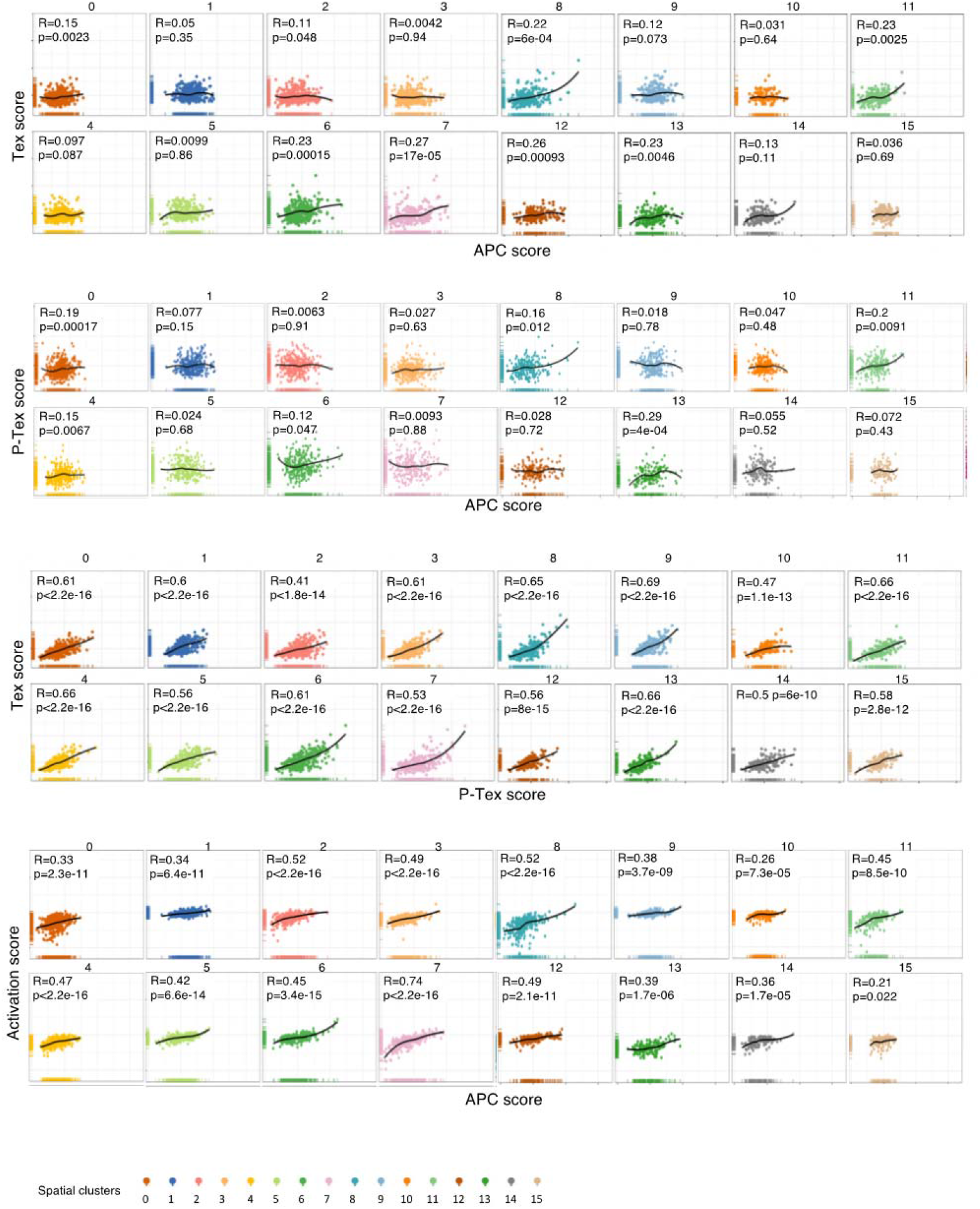
The correlation of Tex, P-Tex and Activation scores with APC scores for each cluster in spatial transcriptomics.

To further confirm the ST results (transcriptomic level) at the proteomic level, we performed multiplex immunofluorescence (mIF) of both canonical APC markers (MHC-II^+^) and selected markers for P-Tex cells (CD8^+^PD1^+^CDK4^+^/MKI67^+^), bulk Tex cells (CD8^+^PD1^+^CDK4^-^MKI67^-^) and bulk CD8^+^ T cells (CD8^+^) on formalin-fixed paraffin embedded (FFPE) tissue originating from HNSCC patients (Fig. 7a-b). Next, we explored average distances of APCs to P-Tex cells, bulk Tex cells and bulk CD8+ T cells (Fig. 7c), respectively. As expected, 90% of the P-Tex cells, bulk Tex and bulk CD8+ cells enriched within a distance of 25um from APCs, forming an intra-tumoral niche(34).

**Figure 7.**
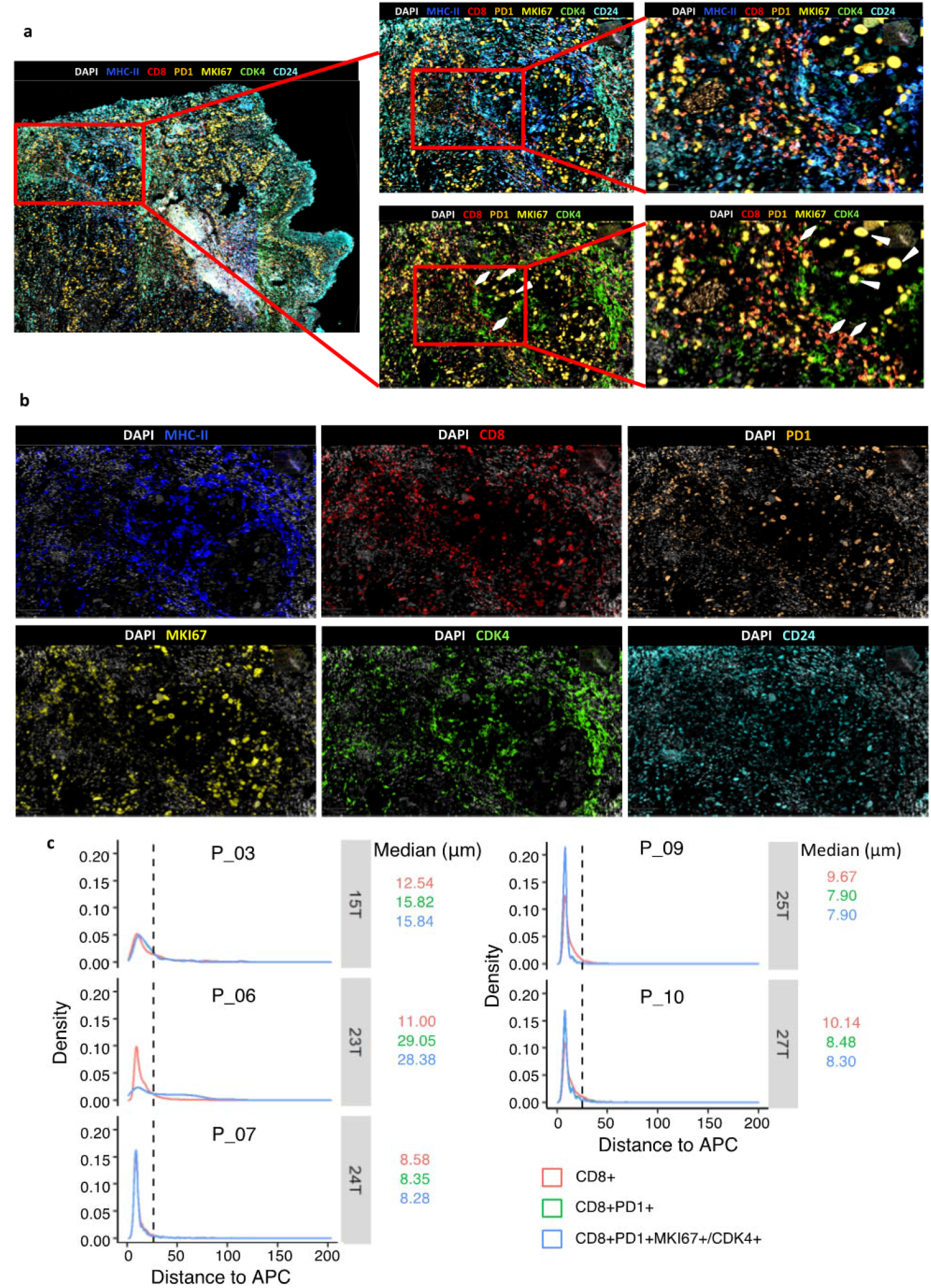
The spatial characteristics of APC, pro-Tex cells and Tex cells in the HNSCC TME. **a-b,** Representative example of HNSCC tumor stained by multiplex IHC, with white triangles and rhombus showing the Texs and P-Tex aggregates in the APC area, respectively, c. Measured distances to APC (MHCÜ+) cells from CD8+T cells, CD8+PD1 +T cells (Tex cells) or CD8+PD1+MKI67+/CDK4+T cells (P-Tex cells) in five representative samples. The dashed lines represent the cutoff distance of 25μm, which indicate that 90% of CD8+T, Tex or P-Tex cells are enriched within a distance of 25um from APCs.

Taken together, P-Tex cells were enriched in the APC aggregation areas, and the signal pathways related to T cell activation and proliferation were activated in these areas, indicating that P-Tex might act as a specific T cell pool that provides modest but persistent effects through its interactions with APCs.

## Discussion

Our current study provided a comprehensive multi-omics characterization of over 49,000 tumor-infiltrating CD3^+^ T cells in HNSCC patients. A special novel P-Tex cluster that expressed high levels of proliferation- and cell cycle-related genes as well as cytotoxic and checkpoint molecules was identified. Our results showed that HPV-positive HNSCC patients who had higher proportions of P-Tex cells had a better survival prognosis. Unexpectedly, we also found that P-Tex cells expressed CDK4 genes as high as cancer cells, which could be simultaneously inhibited by the CDK4 inhibitors. Therefore, we speculated it might be a potential reason for the ineffectiveness of CDK4 inhibitors in treating HPV-positive HNSCC. Furthermore, P-Tex cells were fund to be aggregated in the APC areas where their T cell activation and proliferation signaling pathways were activated. Together, our findings reveal a promising role for P-Tex cells in the prognosis of HPV-positive HNSCC patients by providing modest but persistent anti-tumor effects.

There is accumulating evidences showing that heterogeneity is a hallmark of T cell exhaustion, and a typical three-stage differentiation trajectory (progenitor-transitional-terminal) has been established to depict the corresponding spatiotemporally sequential alterations of gene signatures, functional characteristics as well as epigenetic modifications(9,35). Despite several previous studies have identified similar proliferation Tex clusters in chronic LCMV-infected mouse models using scRNA-seq, little attention was paid to their potential roles in anti-tumor immunity(8,22,36). In this sutudy, we systematically investigated the functional characteristics and developmental trajectory of P-Tex cells by comparing with other CD8^+^ T cell clusters. Our results suggested that P-Tex was an independent branch of Tex cells and might act as a T cell pool, providing modest but persistent anti-tumor immunity through its highly specialized self-renewal and cytotoxic capacity. However, this beneficial on long-term survival outcomes was only observed in HPV-positive HNSCC who had higher proportion of P-Tex.

CDK4/6 inhibitors (e.g., palbociclib, ribociclib and abemaciclib) are promising drugs for various cancers(37), working by specifically inhibiting CDK4/6 proteins, blocking the transition from the G1 to the S phase of the cell cycle and preventing cancer cell progression(38). Notably, *CDK4*, which was highly expressed in cancer cells, was also found to be highly expressed in the P-Tex cells. Our in vitro results showed that CDK4 inhibitors could simultaneously inhibit the cell viability of both cancer cells and P-Tex cells. Due to the fact that P-Tex cells was benefit to the prognosis of HPV-positive HNSCC patients, we speculated that the inhibition of CDK4 inhibitors on P-Tex cells might be one of the reasons why promising treatment outcomes of CDK4 inhibitors were not observed in HPV-positive HNSCC patients(17–19).

Overall, a novel promising P-Tex cluster, which was mainly identified in APC areas of TME, was beneficial to the survival prognosis of HPV-positive HNSCC,. Besides, the inhibitory effect of CDK4 inhibitors on P-Tex cells helps clarify its ineffectiveness in HPV-positive HNSCC patients.

## Materials and Methods

### Ethical statement

This study was conducted in accordance with the Declaration of Helsinki (as revised in 2013) and was approved by the Biomedical Research Ethics Committee of West China Hospital (2021-908), with the individual consent for each participant.

### Specimen collection and processing

Patient’s information was summarized in Table S1. HNSCC tumor tissues and paired adjacent normal tissues were collected during the surgery. Then the tissues were rinsed by 1X PBS, with surrounding necrotic areas being carefully removed and were cut into small pieces of 2–4 mm and reserved in the mixture of 1X DMEM medium (Gibco) and Penicillin-Streptomycin solution (Hyclone). The remaining tissues were fixed into formalin fixed paraffin-embedded blocks (FFPE) for HE staining and multiplex immunofluorescence.

### Preparation of single cell suspensions

The tissue pieces were rapidly transferred into the gentleMACS C Tube containing Human Tumor Dissociation Kit (Miltenyi Biotec, #130-095-929) according to the manufacturer’s recommendation. The dissociated cells were filtered through 40-μm cell strainers to remove clumps. Cell pellets were resuspended in binding buffer after centrifuged and sorted via human CD3 MicroBeads (Miltenyi Biotec, #130-050-101) according to the manufacturer’s recommendation (Note: as for the experiment of sc-RNA seq of overall cells, CD3 sorting was not needed). The overall cells and the sorted CD3^+^ T cells were separately resuspended in HBSS (Gibco) plus 0.04% bovine serum albumin (BSA; Sigma-Aldrich) and tested for cell viability. Cell viability >80% was required for subsequent library constructions.

### Library construction and sequencing

Sc-RNA seq was performed using Chromium Single Cell 5’ Gel Bead and Library Construction Kit (10x Genomics, #PN-1000006, PN-1000020) and Single Cell V(D)J Enrichment Kit Human T cell (10x Genomics, #PN-1000005). Reverse transcription, cDNA recovery, cDNA amplification and library construction were performed according to the manufacturer’s protocol. The constructed libraries were sequenced on NovaSeq 6000 (Illumina) with paired-end sequencing and single indexing.

### Quality control and preprocessing of sequencing data

Cell Ranger count (v3.0)(39) was used to process the raw FASTQ files, align the sequencing reads to Ensembl GRCh38 reference genome (http://cf.10xgenomics.com/supp/cell-exp/refdata-cellranger-GRCh38-3.0.0.tar.gz) and exclude background noise to generate a filtered UMI expression matrix for each cell. The package Seurat (v4.0.4)(40) were used to filter cells that were empty droplets or doublets and that have >5% mitochondrial counts. Next, we normalized the expression matrix via “LogNormalize” and log-transform method. Then, we apply a linear transformation to prepare the expression matrix for next step dimensional reduction.

### Unsupervised clustering of cells and uniform manifold approximation and projection (UMAP) visualization

The high cell-to-cell variable features (top 2000) between cells were used as input to perform Principal Component Analysis (PCA) on the scaled matrix. Subsequently, we employed Harmony (v1.0, R package)(41) to integrate multiple samples and the top 30 dimensions were selected for UMAP with the reduction of ‘harmony’.

We performed Seurat to cluster cells using the Louvain algorithm. The previous reported marker genes (Table S3) were used for the cell cluster annotation with gene functional description and gene expression. Nebulosa (v1.3.0, R package)(42) was applied to perform gene kernel density estimation and visualize cell features on UMAP plot.

### Differential expression and analysis of signaling pathways

To characterize the function of defined clusters, we used Seurat to calculate differentially expressed genes (DEGs) among each cluster, identified marker genes as DEGs with adjusted p value <0.05 and put marker genes into clusterProfiler (v4.0.2, R package)(43) to perform Gene Ontology (GO) enrichment analysis (p <0.01) and Gene Set Enrichment Analysis (GSEA) and visualization.

### Transcription factor regulatory network analysis

To predict the gene regulatory network within cell clusters, we used previous selected top 2000 variable features-barcode matrix from scRNA-seq data as input and applied pySCENIC (v0.11.2)(44) to infer the network activity in each cell cluster.

### Cell score (CS) calculation

We applied AddModuleScore function embedded in Seurat to calculate the specific cell scores in different clusters, which was defined as: the average gene expression of specific gene panel in each cluster, subtract the average gene expression of random control gene sets (45)(Table S3). Functional module scores were based on the expression levels of top 30 genes which were highly correlated with GZMB (cytotoxicity score), PDCD1 (exhaustion score) or MKI67 (proliferation), respectively. TCR-depended T cell activation score was calculated based on the activation gene signature(46). Proliferation score was calculated based on the genes enriched in the GO molecular function term of “cell cycle phase transition”. The specific cluster score (P-Tex, Tex and APC score) were calculated based on marker genes of each cluster listed in Table S3.

We assigned cell cycle scores based on the expression of G2/M and S phase marker genes and predicted the classification of each cell in either G2/M, S or G1 phase in the CellCycleScoring function embedded in Seurat.

### Trajectory analyses

To determine the potential development lineages of T cell subclusters, we converted the previous Seurat object into Monocle3 (v1.0.0, R package)(47) object and inferred the trajectory of T cell subclusters at its proper position in pseudotime. Besides, to visualize the major non-linear components of variation across cells, we applied destiny package (v3.1.1, R package)(48) to perform the 3D diffusion maps to compute the diffusion components of each cell type.

### TCR Clonotype analysis

Cell Ranger VDJ pipeline (v6.1.1, 10× Genomics) was used to process the raw TCR sequence data with default augments and align them to the Ensembl GRCh38 reference (https://cf.10xgenomics.com/supp/cell-vdj/refdata-cellranger-vdj-GRCh38-alts-ensembl-5.0.0.tar.gz). We performed scRepertoire (v1.3.2, R package)(49) to integrate the TCR sequence data with mRNA expression data and used absolute frequency of V(D)J genes to define clonotype groups. The total frequency assigned for different extents of clonal expansion were categorized as follows: Hyperexpanded (20 < X ≤ 200), Multiple (2 < X ≤ 20), Double (1 < X ≤ 2), Single (0 < X ≤1).

### Ligand–receptor interactions

To understand communications among tumor cell clusters, we applied CellChat (v1.1.3, R package)(50) to identify the cell-cell signaling links, inferred the cellular communication network and visualized the major ligand-receptor interaction between each cell cluster.

### Library construction of spatial transcriptome

Representative HNSCC tumor samples were collected for the spatial transcriptomic sequencing. Samples were cut into 6.5 x 6.5 mm pieces, embedded in Optimal Cutting Compound (OCT) media and quickly frozen on dry ice. The frozen tissues were cryosectioned at 10-μm thickness by using the Thermo Scientific CryoStar NX50 cryostat and were placed in the capture area frames on the 10x Visium Spatial slides. Each sample slide was stained with H&E (Hematoxylin Dako #S3309, Eosin, Dako #CS701, bluing buffer #CS702) and the brightfield images were captured via Leica whole-slide scanner at 10X resolution.

Following tissue permeabilization, reverse transcription and cDNA amplification were processed by using Reagent Kit (10× Genomics, #PN-1000184, PN-1000193).Visum spatial libraries were constructed using Visum Spatial Library Construction kit (10x Genomics, #PN-1000184) according to the manufacturer’s protocols. Finally, the libraries were sequenced using the Illumina Novaseq6000 at least 100,000 reads per spot via pair-end 150 bp (PE150) reading strategy (performed by CapitalBio Technology, Beijing).

### Functional scoring and visualization of spatial transcriptome data

We performed alignment, filtering, barcode counting, and UMI counting by the Spaceranger count (v1.3.0) to generate feature-barcode matrix. We performed normalization, high-variance features detection (top 2000 genes), dimensionality reduction and clusters identification (resolution = 1.0) for the spatially barcoded gene expression data via the standard Seurat pipeline. The P-Tex, Tex and APC scoring algorithms of spatial transcriptome were similar to the module scoring algorithm of scRNA transcriptome (AddModuleScore) and the gene list of each module was list in Table S5. Co-localization of P-Tex, Tex and APC scores were verified by cor.test (stats, v3.6.2, R package)(51).

### Multiplex immunohistochemistry

Ten formalin-fixed paraffin-embedded tissue (FFPE) of HNSCC tumors were sectioned to 4-μm thick for the subsequent multiplex immunohistochemistry via the OPAL Polaris system (Akoya Biosciences). After deparaffinization and hydration, the FFPE slides were manually stained with the CD8 (clone C8/144B, CST, #70306S), PD-1 (clone D7D5W, CST, #84651T), anti-Ki67 (clone SP6, Abcam, #ab16667), CDK4 (clone D9G3E, CST, #12790), CD24 (10600-1-AP, Proteintech, #10600-1-AP), Anti-EPCAM (clone EPR20532-225, Abcam, #ab223582) and Anti-HLA-DR (clone EPR3692, Abcam, #ab92511) antibodies. The sections were counterstained with spectral DAPI (Akoya Biosciences). The stained slides were imaged and scanned using the Vectra Polaris multispectral imaging system.

### Cell staining strategies for flow cytometry

Single cell suspensions (100μL) of HNSCC tumor tissues were stained with CD3 (BD Pharmingen™, #555332), CD8(CST, #300908), PD1(Biolegend, #329920), UBE2C (Santacruz, #Sc271050), and KLRB1(Biolegend, #339917) antibodies at 4 °C for 30 min under dark conditions. And 7-Aminoactinomycin D (7-AAD) was used for live/dead discrimination. P-Tex cells were defined as 7AAD-CD3^+^CD8^+^PD1^+^ UBE2C^+^cells and T-Tex cells were defined as 7AAD-CD3^+^CD8^+^PD1^+^KLRB1^+^UBE2C^-^ cells.

### In vivo cell function assays

#### Cell culture and proliferation assay

The sorted P-Tex cells and T-Tex cells were cultured in RPMI media containing 10% FBS, penicillin, streptomycin and 20 IU/mL IL-2, and stimulated with T Cell TransAct (Diluted at 1:100, T Cell TransAct™, human, Miltenyi Biotec, #130-111-160), with fresh medium replaced every 3 days. After 14 consecutive days, proliferation rate of cells was assessed by flow cytometry.

#### Self - renew assay

The sorted P-Tex and T-Tex cells were cultured in the RPMI media containing 10%FBS, human IL-2 (20 IU/mL), penicillin and streptomycin in 96-well plates (15,000 cells/well). At the 5^th^, 10^th^ and 15^th^ day of growth, cell counting was performed by flow cytometry. Cell viability was determined by propidium iodide (PI) staining on the 15^th^ and 20^th^ day of cell growth. And on the 9^th^ and 16^th^ day, P-Tex and T-Tex cells were stained by UBE2C (Santacruz, #Sc271050), PD1 (Biolegend, #329920), KLRB1 (Biolegend, #339917) and the protein expression was detected by flow cytometry. Software FlowJo was used for data analysis.

#### CDK4/6 inhibition test

The sorted P-Tex and T-Tex cells (50000 cells/well) and the cancer cell line (FD-LSC-1, 4000 cells/well, donated by State Key Laboratory of Biotherapy, West China Medical School, Sichuan University)(52) were transferred into 96-well plates, treated with Abemaciclib in gradient concentrations (0nM, 10nM, 100nM and 500nM) for 24h, 48h and 72h, respectively. Cell proliferation was detected by Cell Counting Kit-8 (CCK-8) according to the manufacture’s instruction.

### Survival analysis

We further analyzed the transcriptome data of 500 HNSCC tumor samples (HPV negative: n=410; HPV positive: n=90) in TCGA cohort(53). Survival analysis related to gene expression level and functional module score in different HPV status was conducted through the Tumor Immune Estimation Resource (TIMER; cistrome.shinyapps.io/timer)(54). Besides, to predict the proportion of P-Tex, Teff, and T-Tex cells in HPV positive and HPV negative HNSCC TCGA cohort, we used CIBERSORT software to deconvolve our scRNA seq data (534 specific marker genes, Table S3) into the TCGA bulk transcription data for clustering.

### Statistical analysis

Statistical analysis was performed using R (Version 3.6.3). Wilcoxon rank-sum tests and Chi-square tests were used to compare variables. The hazard ratio (HR) and survival curves was estimated via a Cox regression model. A statistical significance was considered at P < 0.05.

### Data availability statement

Sequencing data have been deposited in GSA for human under accession codes PRJCA012438.

## Acknowledgments

We would like to thank the staff and students in the Department of Oto-Rhino-Laryngology, West China Biomedical Big Data Center, and Research Core Facility of West China Hospital for giving us kind support of sample collection and experiments.

## Author contributions

Conceptualization: RJJ, ZY, YHP, ZW, CF

Methodology: YHP, CDN, RYF, MMZ, LL, WY, ZY, CXM, WSS, LJ

Investigation: QK, SY, CJR, YXW, SXL, HSH, YSS, WHY, PXC, LDB, YL

Visualization: CL, YZY, ZYB, ZMJ, YBW, ZXX

Supervision: XW, LG

Writing—original draft: RJJ, CDN, QK, RYF, MMZ

Writing—review & editing: RJJ, ZY, YHP, ZW, CF, XW, LG

## Competing interests

Authors declare that they have no competing interests.

## Supplementary Tables

**Supplementary Table 1**: Patients information.

**Supplementary Table 2**: Cell numbers of single T cells from 14 HNSCC samples by scRNA-seq.

**Supplementary Table 3**: Marker genes of different cell clusters applied in scRNA-seq.

**Supplementary Table 4**: Transcriptional regulators of top expressed genes in each T cell clusters.

**Supplementary Table 5**: Functional cell scores of each cell cluster.

**Supplementary Table 6**: Numbers of shared clonotypes among each cell clusters.

**Supplementary Table 7**: The overlap coefficients among each cluster.

**Supplementary Table 8**: Cell numbers of different HNSCC TME subclusters by scRNA-seq.

**Supplementary Table 9**: The proportion of P-Texs, T-Tex and Teff clusters in HPV^+^ and HPV^-^ HNSCC samples in TCGA database.

**Supplementary Table 10**: The communication network of signaling pathways among different cell clusters derived by ligand–receptor interactions.

**Supplementary Table 11**: The P-Tex, Tex and APC scores in spatial transcriptomic.

**Supplementary Table 12**: The key resources.

## Notes

**Conflicts of interest**, The authors declare no potential conflicts of interest.

### Competing Interest Statement

The authors have declared no competing interest.

